# Postsynaptic adaptations in direct pathway muscarinic M4-receptor signaling follow the temporal and regional pattern of dopaminergic degeneration

**DOI:** 10.1101/2025.03.03.641233

**Authors:** Beatriz E. Nielsen, Christopher P. Ford

**Author notes:** **Lead / Corresponding author:** Christopher Ford **Address for Correspondence:** Department of Pharmacology; University of Colorado School of Medicine, 12800 E 19^th^ Ave, Aurora, CO, 80045, USA.

## Abstract

In Parkinson’s disease (PD), imbalances in dorsal striatum (DSt) output pathways leading to motor dysfunction are thought to be driven by the loss of dopamine (DA) itself and the disruption of its coordinated modulation with acetylcholine (ACh). While the gradual decline of DA across striatal regions over time is a defining characteristic of PD, less is known about the adaptive and/or pathological alterations in cholinergic signaling that develop throughout disease progression in response to DA loss. Here, we examined changes in cholinergic modulation of striatal direct pathway medium spiny neurons (dMSNs) in mice that were partially or completely depleted of DA, in order to model early and advanced stages of PD. We found a reduction in muscarinic M4 receptor signaling that began in the dorsolateral striatum (DLS) following a partial loss of DA, yet was not evident in the dorsomedial region (DMS) until the dopaminergic lesion was nearly complete. Combining electrophysiological, pharmacological and 2-photon imaging approaches, we determined that this decrease was the result of reduced postsynaptic M4 receptor function, and was not accounted for by changes in ACh release or clearance. Replacing the partial loss of endogenous DA with levodopa could not rescue the dysfunctional M4 receptors. Together, these findings reveal how changes in cholinergic modulation closely follow the temporal and regional pattern of dopaminergic degeneration, which is critical for understanding their shared role in PD progression, and for developing alternative therapeutic interventions.

## INTRODUCTION

Parkinson’s disease (PD) is a neurodegenerative disorder characterized by the progressive loss of substantia nigra pars compacta (SNc) dopamine (DA) neurons and their projections to the dorsal striatum (DSt), which leads to motor symptoms such as bradykinesia, tremor, and rigidity. Medium spiny neurons (MSNs), which are the output cells of the striatum, are a major target of dopaminergic modulation and form two efferent pathways that differ in their expression of DA receptors: D1-expressing direct pathway MSNs (dMSNs) and D2-expressing indirect pathway MSNs (iMSNs). Motor impairment in PD is thought to result from an imbalance in striatal output circuitry following the loss of DA modulation^1,2^ and thus, DA replacement with its precursor levodopa (L-DOPA) remains the gold-standard treatment^3^.

In addition to DA, MSNs are also modulated by acetylcholine (ACh), which arises from cholinergic interneurons (ChIs) and interacts with DA to regulate striatal motor output^4,5^. Alterations in ChI excitability, ACh levels and cholinergic signaling have been identified in different animal models to contribute to parkinsonian motor deficits^6–17^, positioning the cholinergic system as a promising non-dopaminergic therapeutic target. However, despite this central role and therapeutic potential, it remains unclear how pathological and/or compensatory changes in striatal cholinergic system develop as the disease progresses over time and space.

Multiple adaptive changes have been found to arise in striatal circuits following DA loss, which are thought to be compensatory and beneficial at initial stages of degeneration yet may become deleterious over the long-term or when conventional medications are initiated^18^. Such is the case for reduced cholinergic modulation of dMSNs through Gα_i/o_-coupled M4 muscarinic receptors in response to DA depletion, which might be an initial compensation for diminished Gα_olf/s_-coupled D1 receptor signaling that eventually contributes to parkinsonian and L-DOPA-induced dyskinetic motor deficits^17^. Although the development of compensatory mechanisms may delay the onset of motor symptoms^18^, little is known about changes in M4-mediated modulation of the direct pathway at early stages of dopaminergic degeneration, when striatal DA is only partially decreased.

The progressive decline in DA in PD follows a characteristic regional pattern across DSt regions which is thought to be the result of selective vulnerability of different SNc subpopulations^19–23^. The earliest and most severe loss occurs in the ventral tier of the SNc, which projects to the putamen or dorsolateral striatum (DLS), whose associated functions, such as locomotion and habitual actions, are typically impaired in PD^24^. In contrast, the dorsal tier of the SNc projects to the caudate nucleus or dorsomedial striatum (DMS), which is involved in goal-directed behaviors and remains relatively spared^24^. While we have previously reported that direct pathway M4-mediated cholinergic signaling was reduced in the DLS following a complete DA lesion^17^, it remains unknown whether this alteration also occurs at the DMS, where dopaminergic denervation is incomplete. To address this, we compared alterations in M4-signaling in dMSNs across DLS and DMS when mice were partially or completely depleted of DA. We found that reductions in M4-signaling began in the DLS with mild DA lesions and progressed to the DMS only with complete DA loss. As M4 receptor constitutes a promising therapeutic target, unveiling the temporal and regional pattern of its alterations is critical for understanding PD pathophysiology and the advent of novel therapies.

## MATERIALS AND METHODS

### Animals

Animal experiments were conducted according to the protocols approved by Institutional Animal Care and Use Committee (IACUC) at the University of Colorado School of Medicine. Animals were group-housed in a temperature- and humidity-controlled environment on a 12 h light-dark cycle, with water and food available *ad libitum*. All behavioral experiments were conducted during the light phase. Both adult male and female 10–11-week-old wildtype C57BL/6J (WT) (Jackson Laboratory, RRID:IMSR_JAX:000664) and Drd1-Cre heterozygote (D1-Cre^+/-^) (MMRRC, RRID:MMRRC_034258-UCD) mice were used.

### Stereotaxic surgery

Mice (7-8 weeks old) were initially anesthetized with isoflurane in order to be mounted in a stereotaxic frame (Kopf Instruments), where they were kept under constant 2% isoflurane anesthesia during the entire procedure. AAV viruses were unilaterally injected using a Nanoject III (Drummond Scientific) at 2 nL/s into the center of the DSt, at the following coordinates relative to bregma (in mm): AP +0.9, ML ± 1.85, DV −2.9. The needle was maintained in the target site for 5 min to allow virus diffusion. For experiments involving conditional GIRK2 expression in D1-expressing cells, 400 nL of AAV9.hSyn.DIO.tdTomato.T2A.GIRK2 (University of Pennsylvania Viral Core, RRID: SCR_022432, V5688R) was injected in D1-Cre mice. For experiments requiring non-selective striatal expression of GRAB_ACh 3.0_ sensor, WT mice were injected with 400 nL of AAV9-hSyn-ACh4.3 (WZ Biosciences, RRID:Addgene_121922). For more details, see dx.doi.org/10.17504/protocols.io.kqdg3x9j1g25/v1.

### Unilateral 6-OHDA lesion model of PD

6-OHDA (6-hydroxydopamine hydrobromide, Sigma-Aldrich, 162957) was dissolved in sterile saline and injected unilaterally (1 µL) either at a low dose (1 µg/µL, 6-OHDA LD) or a high dose (4 µg/µL, 6-OHDA HD) into the mouse medial forebrain bundle (MFB) during stereotaxic surgery at the following coordinates relative to bregma (in mm): AP - 1.2, ML +1.3, DV −4.75. Control or sham mice received sterile saline instead (1 µL). At least 30 min prior to 6-OHDA injections, mice were pretreated (i.p.) with 25 mg/kg desipramine and 5 mg/kg pargyline dissolved in sterile saline. To minimize mortality following DA depletion, special care was conducted during at least 1-2 weeks post-surgery. Mice remained in a heat pad and received daily sterile saline i.p. injections, food pellets soaked in water, sunflower seeds and nutritionally fortified water gel (DietGel Recovery, ClearH20) as needed. For more details, see dx.doi.org/10.17504/protocols.io.81wgbxyjylpk/v1.

### Levodopa treatment

Three weeks after 6-OHDA LD injection, a subset of mice was examined by the cylinder test to confirm a partial loss of DA (see Cylinder test). Mice then started receiving daily i.p. injections of 2 mg/kg of L-DOPA (3,4-dihydroxy-L-phenylalanine) (Sigma-Aldrich, D9628) plus 12 mg/kg of benserazide hydrochloride (Sigma-Aldrich, B7283) dissolved in sterile saline for 7-8 days, as previously described^17^. Brain slices for electrophysiology recordings were obtained 30 min – 1 h after the last L-DOPA administration. For more details, see dx.doi.org/10.17504/protocols.io.4r3l29dpjv1y/v1.

### Brain slices preparation

Mice were anesthetized with isoflurane and transcardially perfused with ice-cold sucrose cutting solution containing (in mM): 75 NaCl, 2.5 KCl, 6 MgCl_2_, 0.1 CaCl_2_, 1.2 NaH_2_PO_4_, 25 NaHCO_3_, 2.5 D-glucose and 50 sucrose, bubbled with 95% O_2_ and 5% CO_2_. Coronal striatal slices (240 µm) were cut in the same ice-cold sucrose cutting solution. Slices were incubated for at least 45 min at 32°C in artificial cerebro-spinal fluid (aCSF) containing (in mM): 126 NaCl, 2.5 KCl, 1.2 MgCl_2_, 2.5 CaCl_2_, 1.2 NaH_2_PO_4_, 21.4 NaHCO_3_, and 11.1 D-glucose, bubbled with 95% O_2_ and 5% CO_2_. To prevent excitotoxicity, 10 µM MK-801 was added. After incubation, slices were transferred into a recording chamber and constantly perfused with aCSF warmed to 32 ± 2°C at a rate of 2 mL/min. Neurons were visualized using a BX51WI microscope (Olympus) with infrared and green LEDs (Thorlabs). For more details, see dx.doi.org/10.17504/protocols.io.4r3l2omjpv1y/v1.

### Ex vivo electrophysiology

All recordings were conducted in the DSt using Axopatch 200B amplifiers (Molecular Devices). Patch pipettes (1.5 – 2 MΩ) (World Precision Instruments) were made using a pipette puller (Narishige, PC-10). Pipettes for voltage-clamp whole-cell recordings contained (in mM): 135 D-Gluconate (K), 10 HEPES(K), 0.1 CaCl_2_, 2 MgCl_2_ and 10 BAPTA-tetra potassium, with 1 mg/mL ATP, 0.1 mg/mL GTP, and 1.5 mg/mL phosphocreatine (pH 7.35, 275 mOsm). Cells were held at −60 mV. No series resistance compensation was applied, and cells were discarded if series resistance was ≥15 MΩ. All recordings from GIRK2^+^ dMSNs were performed in regions showing robust reporter fluorescence to ensure less variability of GIRK2 outward currents between cells and mice^17,25,26^. Endogenous ACh release was triggered by electrical stimulation (0.5 ms) using a monopolar glass stimulating electrode filled with aCSF, positioned consistently 200 µm away from the recorded cell. Recordings were acquired with Axograph 1.76 (Axograph Scientific; RRID SCR_014284) at 10 kHz and filtered to 2 kHz, or with LabChart (ADInstruments; RRID:SCR_017551) at 1 kHz. For more details, see dx.doi.org/10.17504/protocols.io.j8nlkooowv5r/v1. All drugs were bath applied and recordings were performed in aCSF containing 10 µM DNQX, 100 µM picrotoxin, 300 nM CGP 55845, 10 mM SCH 23390, 10 µM DHβE and 200 nM sulpiride to isolate muscarinic cholinergic transmission.

### 2-photon imaging GRAB_ACh 3.0_ recordings

2-photon imaging was performed using a 2-photon laser scanning microscopy system, custom-built on a BX51WI microscope (Olympus). A Ti:Sapphire laser (Chameleon Ultra I; Coherent) was tuned to emit pulsed excitation at 920 nm and scanned using a pair of X-Y galvanometer mirrors (6215, Cambridge Technology). Emitted fluorescence was collected through a water-immersion objective (60X, Olympus), a dichroic mirror (T700LPXXR, Chroma) and filters (ET680sp and ET525/50 m-2P, Chroma), and was detected using a GaAsP photomultiplier tube (PMT, H10770PA-40, Hamamatsu). A current preamplifier (SR570, Stanford Research Systems) was used to convert the output to voltage, which was then digitized by a data acquisition card (PCI-6110, National Instruments). ACh release was triggered by electrical stimulation (0.5 ms) with the electrode positioned in the center of square regions of interest (25 µm x 25 µm) and peak fluorescence changes of GRAB_ACh3.0_ were measured using a custom software (Toronado; https://github.com/StrowbridgeLab/Toronado-Laser-Scanning) under two acquisition modes, rasterized frame scans or 2-photon non-raster point scanning as previously described^27^. For the former, the mean frame was quantified using Fiji (ImageJ, RRID:SCR_002285). For the latter, the laser was repeatedly scanned across a small circular path (150 nm diameter) at a selected spot of interest inside the square region and fluorescence was continuously collected from that spot. The PMT signal was converted by the same preamplifier (SR570, Stanford Research Systems; sensitivity 100 nA/V), but further filtered to 500 Hz with the gain increased two-fold (FLA-01, Cygnus Technologies). Then the signal was acquired using a data acquisition device (ITC-18, HEKA Instruments) and recorded and analyzed using Axograph X (Axograph Scientific; RRID SCR_014284). For more details, see dx.doi.org/10.17504/protocols.io.kxygx33yog8j/v1.

### Immunohistochemistry and fluorescence imaging

Anesthetized mice were perfused transcardially with cold PBS followed by cold 4% paraformaldehyde in PBS, pH 7.4. Brains were post-fixed in 4% PFA at 4°C for an additional period of 2 h, equilibrated in 30% sucrose solution for 48 h, and rapidly frozen in embedding freezing medium (Fisher Scientific). DSt and/or midbrain coronal slices of 30 µm thickness were obtained using a Leica CM1950 cryostat (Leica Microsystems). For TH-immunohistochemistry, sections were mounted on slides and blocked in 5% normal donkey serum in PBS-T (0.3% Triton X-100) for 1 h at RT. Slides were then incubated with rabbit anti-TH primary antibody (1:200, Millipore AB152, RRID: AB_390204) overnight at 4°C. After PBS washes, slides were incubated with donkey anti-rabbit Alexa Fluor 488 secondary antibody (1:500, Life Technologies A21206, RRID: AB_2535792) for 1 h at RT and washed afterward with PBS.

For immunostaining GIRK2 and choline acetyltransferase (ChAT), free floating DSt coronal sections in a multiwell plate were blocked in 10% normal donkey serum in 1% PBS-T for 1 h at RT. Slices were then incubated with rabbit anti-GIRK2 (1:500, Alomone labs APC-006, RRID: AB_2040115) and goat anti-ChAT (1:400, Milipore AB144P, RRID: AB_2079751) primary antibodies overnight at 4°C on shaker. After washes with 0.5% PBS-T, slices were incubated with donkey anti-rabbit Alexa Fluor 488 (1:500, Life Technologies A21206, RRID: AB_2535792) and donkey anti-goat Alexa Fluor 647 (1:500, Abcam ab150131, RRID:AB_2732857) secondary antibodies for 1 h at RT on shaker and washed afterward with PBS. Following immunostaining, sections were finally mounted with an anti-fading mounting media containing DAPI (Vectashield, Vector laboratories). Fluorescent images were acquired using a slide scanner (VS120, Olympus) or a confocal microscope (Zeiss LSM780, Carl Zeiss) and processed in Fiji (ImageJ, RRID: SCR_002285). For more details, see dx.doi.org/10.17504/protocols.io.14egn7yyzv5d/v1.

### Chemicals

Picrotoxin, MK-801, DNQX, DHβE, SCH 23390, sulpiride, acetylcholine, oxotremorine M, and ambenonium were purchased from Tocris Bioscience. Desipramine, 6-OHDA, L-DOPA and benserazide were from Sigma-Aldrich. CGP55845 was obtained from Hello Bio, BAPTA from Invitrogen and pargyline from Abcam.

### Cylinder test

Forelimb use asymmetry was assessed by cylinder test at 3 weeks following stereotaxic injections. Mice were habituated to the behavioral room for 40 min prior to the test. Individual mice were then placed in a clear plastic cylinder (10.5 cm diameter; 14.5 cm height) with mirrors behind for complete vision and were video recorded during 5 min for later post-hoc scoring. Only wall contacts executed with fully extended digits were scored. Data was expressed as a percentage of touches performed with the forelimb contralateral to the injected side with respect to the total paw use (ipsilateral + contralateral). Forelimb lateralization revealed by this test was conducted to verify the extent of DA-depletion in parkinsonian mice prior electrophysiology and imaging experiments. Only mice whose contralateral paw use was lower than 45% and 40% for 6-OHDA LD and HD conditions respectively were included for experiments. For more details, see “Cylinder test: forelimb asymmetry” at dx.doi.org/10.17504/protocols.io.j8nlkoon6v5r/v1.

### Rotarod

Motor balance and coordination were assessed by the rotarod test at 3 weeks following stereotaxic injections, using an accelerated protocol from 3 to 30 rpm in 5 min (Med Associates). Mice were habituated to the behavioral room for 40 min prior to the test. Each mouse underwent 3 trials on the same day with a separation of 10 min between trials, without previous training. The latency to fall or time to reach maximum speed was recorded, and the data was expressed as the average of the two best trials. For more details, see “Rotarod: balance and coordination” at dx.doi.org/10.17504/protocols.io.j8nlkoon6v5r/v1.

### Experimental Design and Statistical Analysis

Experiments were conducted in both male and female adult mice and the collected data was analyzed using Prism 10 (GraphPad 10.0.2, RRID:SCR_00306). Data sets that passed the Shapiro-Wilk test for normality were analyzed using parametric tests, otherwise, non-parametric tests were applied. For comparison between two groups, the following two-tailed statistical tests were conducted as appropriate: unpaired or paired Student t-test, Mann-Whitney U test or Wilcoxon matched-pairs signed rank test. For comparisons between more than two groups, the following statistical tests were applied as appropriate: One-way analysis of variance (ANOVA) or Kruskal-Wallis, Two Way ANOVA or Mixed-model ANOVA with Geisser-Greenhouse’s correction. Repeated measures (RM) version of those tests were performed for matched or paired data. When significant differences were found in ANOVA tests, Tukey’s, Dunn’s or Holm-Sidak’s post-hoc multiple comparison tests were performed respectively. Statistical significance was established as p≥0.05 (n.s.), p<0.05 (*), p<0.01 (**), p<0.001 (***), and p<0.0001 (****). Summary data (mean ± SEM), exact p-values, n (number of cells), N (number of animals), and specific statistical tests and comparisons for each experiment are stated in figure legends and Supplemental Table S1 in an extended format.

## RESULTS

### Partial DA lesion replicates parkinsonian-like uneven pattern of dopaminergic denervation across dorsal striatum regions

To mimic and compare early and advanced stages of parkinsonism, we induced partial or near complete dopaminergic lesions by unilaterally injecting 6-hydroxydopamine (6-OHDA, 1 µL) into the medial forebrain bundle (MFB) of mice at either a low (1 µg/µL; 6-OHDA LD) or a high (4 µg/µL; 6-OHDA HD) dose^28–30^ (Figure 1A). Control mice were injected with saline (Figure 1A). Three to four weeks after 6-OHDA injection, tyrosine hydroxylase (TH) immunoreactivity was reduced in the DSt due to the loss of dopaminergic fibers, and in the SNc due to the loss of dopaminergic neurons (Figure 1B-1C). While the high-dose of 6-OHDA resulted in a ∼90-95% reduction in total TH-immunoreactivity in DSt and SNc relative to sham vehicle-injected hemispheres^17,28,30^, the low-dose induced a graded lesion with a ∼50-60% decrease (Figure 1B-1C). This reduction correlates with the estimated ∼50% reduction in putamen DA fibers required to produce clinical motor symptoms in PD^22^, and is similar to the ∼57% reduction in evoked DA release we have previously reported with low-dose 6-OHDA partial lesions^29^.

**Figure 1:**
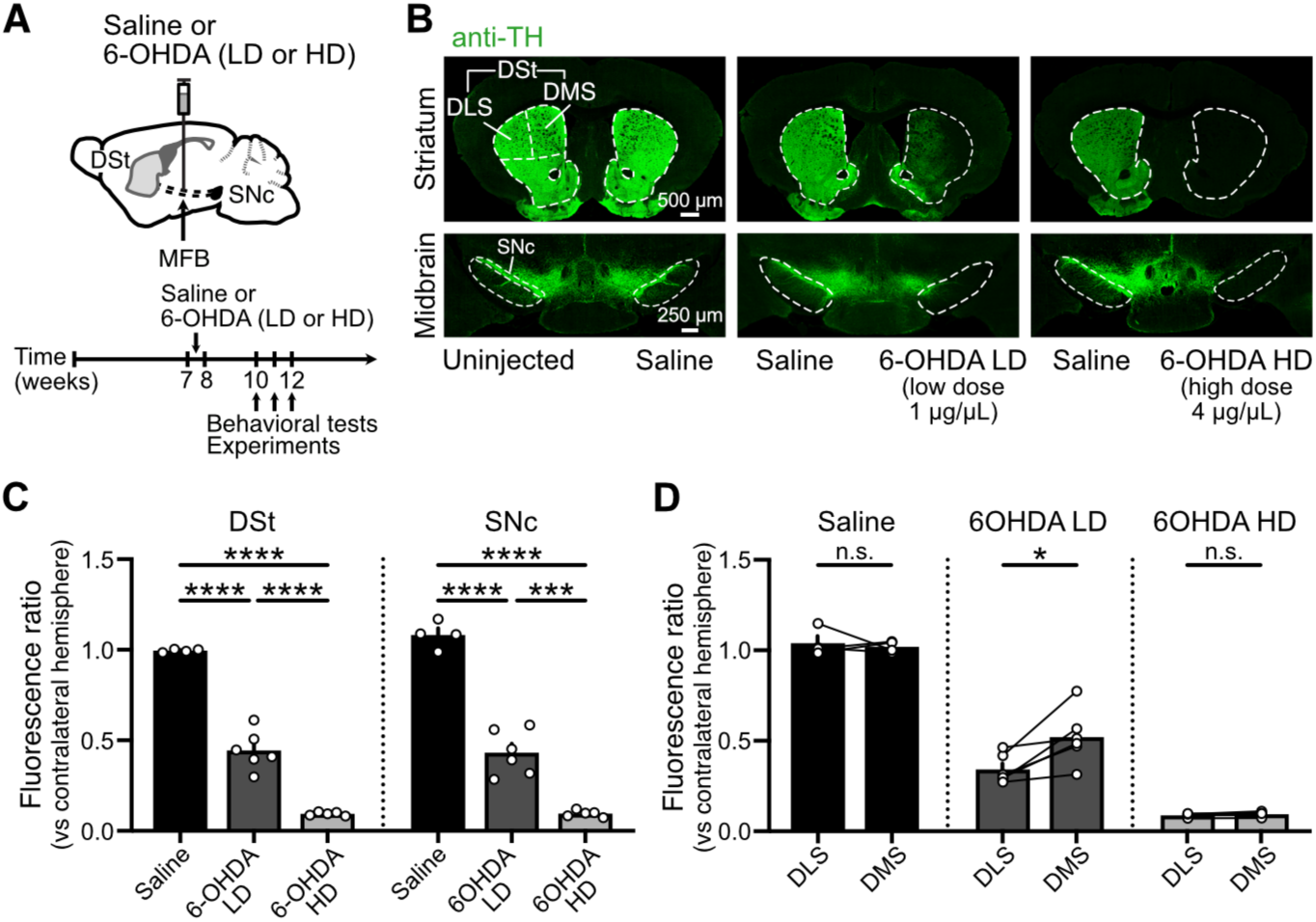
Partial DA depletion by a low-dose of 6-OHDA leads to differential dopaminergic denervation across striatal regions. **(A)** Schematics of saline or 6-OHDA injections into the MFB (top) and timeline for experiments (bottom). **(B)** Striatal and midbrain coronal sections showing TH-immunoreactivity following unilateral injections of 1 µL of saline, 6-OHDA LD (1 µg/µL) or 6-OHDA HD (4 µg/µL) into the MFB. **(C)** Quantification of TH-immunoreactivity in DSt and SNc expressed as the fluorescence ratio of the injected hemisphere vs the uninjected/saline-injected hemisphere (Saline N=4, 6-OHDA LD N=6, 6-OHDA HD N=5; One-Way ANOVA p<0.0001, Tukey’s post-hoc test). **(D)** Quantification of TH-immunoreactivity across DMS and DLS for each group (Saline N=4, paired t-test p=0.7013; 6-OHDA LD N=6, paired t-test p=0.0165; 6-OHDA HD N=5, paired t-test p=0.1327). Summary data is mean ± SEM. Extended statistical data is provided in Supplemental Table S1. n: number of cells, N: number of mice; n.s. p>0.05; *p<0.05; **p<0.01; ***p<0.001; ****p<0.0001.

As the differential sensitivity of dopaminergic neurons to degeneration leads to uneven striatal DA depletion in PD, we further dissected the extent of 6-OHDA-induced DA lesion across striatal regions in our animal model. In accordance with earlier stages of neurodegeneration, the low-dose of 6-OHDA resulted in a greater loss of TH-immunoreactivity in the DLS (∼66%) compared to the DMS (∼50%), while the high-dose of 6-OHDA led to a near complete depletion of DA in both striatal regions, mimicking more advanced stages of the disease (Figure 1B, 1D).

The degree of DA denervation in the putamen has been considered an indicator of the severity of motor symptoms in PD patients^31^. Thus, we conducted a correlation analysis between the extent of dopaminergic fiber loss across striatal regions in our rodent model with the motor performance in two behavioral assays: the cylinder and the rotarod tests (Figure 2A). These assays were aimed to reveal the asymmetry in forelimb use and the overall impairment in motor balance and coordination in parkinsonian mice. We found that the severity of motor deficits in both tests was significantly correlated with the loss of TH^+^-fibers in the DSt as a whole, as well as when examining region-specific dopaminergic denervation in the DLS and DMS (Figure 2A). The presence of motor impairment allowed us to distinguish partial- and total-lesioned mice from sham saline-treated control mice (Figure 2B-2C) and was subsequently used as a criteria for successful dopaminergic lesions and the inclusion of those animals in the parkinsonian experimental groups. Thus, a more moderate DA lesion reproduces the parkinsonian-like uneven pattern of dopaminergic fiber denervation in the DSt, providing a model to examine the adaptations in M4-mediated cholinergic signaling at early versus advanced stages of neurodegeneration, as well as across DMS and DLS regions.

**Figure 2:**
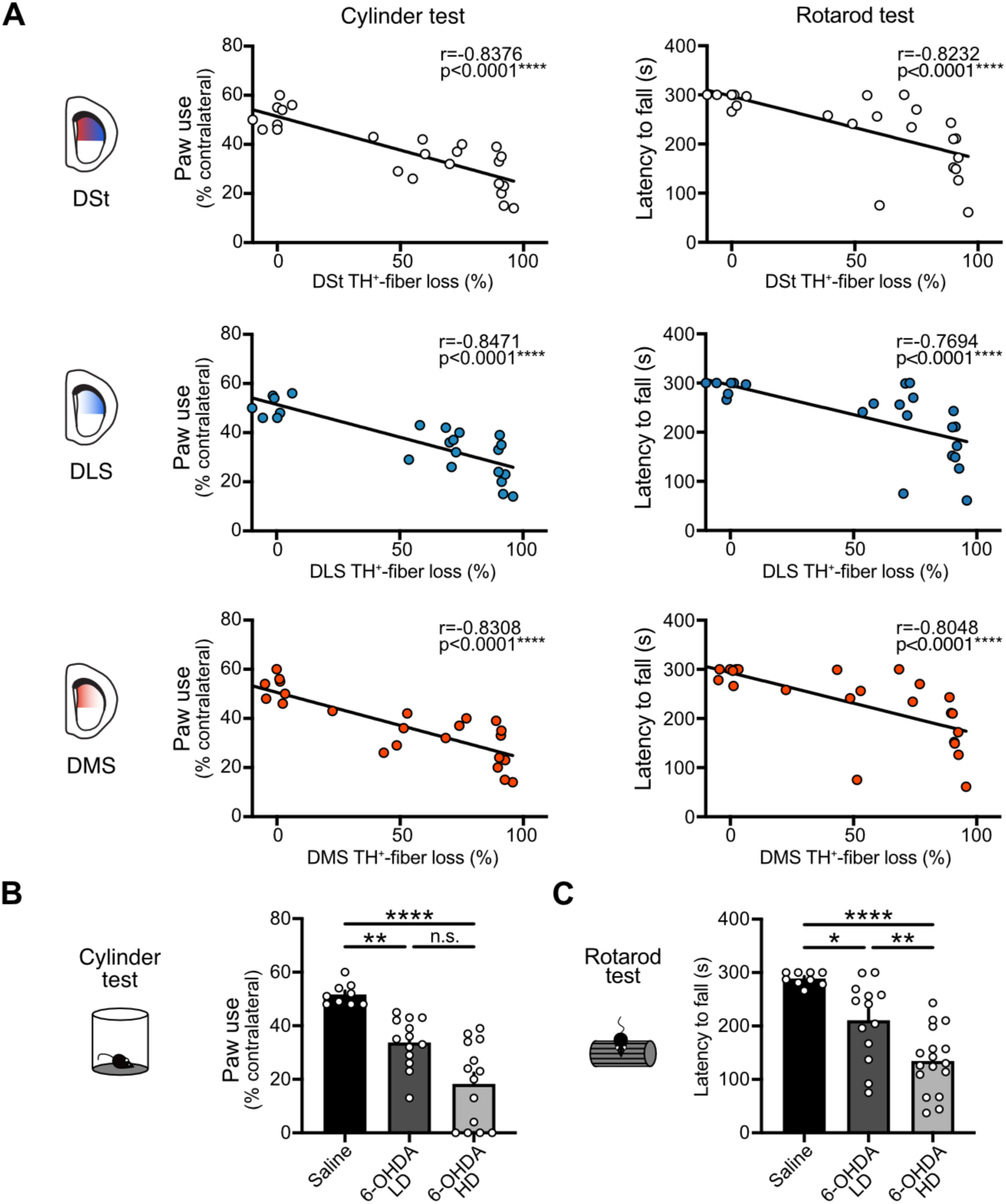
Dopaminergic lesions induced by different doses of 6-OHDA correlate with the severity of motor impairment. **(A)** Correlation of cylinder test (left) and rotarod (right) performances with TH^+^-fiber loss in the entire DSt, DLS or DMS (N=24; Pearson correlation for cylinder test, Spearman correlation for rotarod test). **(B)** Summary data of contralateral paw use in cylinder test (Saline N=9, 6-OHDA LD N=13 and 6-OHDA HD N=14; Kruskal-Wallis p<0.0001; Dunn’s post-hoc test). **(C)** Summary data of rotarod test (Saline N=9, 6-OHDA LD N=13 and 6-OHDA HD N=14; One-Way ANOVA p<0.0001; Tukey’s post-hoc test). Summary data is mean ± SEM. Extended statistical data is provided in Supplemental Table S1. n: number of cells, N: number of mice; n.s. p>0.05; *p<0.05; **p<0.01; ***p<0.001; ****p<0.0001.

### Partial and complete DA depletion differentially reduce M4-mediated cholinergic signaling across DSt regions

Following a near complete depletion of DA in the DLS, the strength of M4-mediated cholinergic transmission in dMSNs is reduced^17^. We wanted to examine whether this adaptation occurs at early stages of dopaminergic degeneration and across DSt regions. As above, we induced partial and total dopaminergic lesions by unilaterally injecting either the low- or the high-dose of 6-OHDA into the MFB (Figure 3A). To measure direct pathway M4-mediated cholinergic signaling and given the lack of endogenous conductances efficiently coupling to M4 receptors, we selectively expressed exogenous G-protein-coupled inwardly rectifying K^+^ channels (GIRK2; Kir3.2) in dMSNs, where GIRK2 can couple to M4 receptors and the resulting potassium current provides an electrophysiological readout for receptor activation^17,32,33^ (Figure 3A). This approach, although involving an exogenous effector, has been extensively validated and provides a robust and reliable measurement of M4 receptor activation, with temporal and spatial scales comparable to endogenous signaling and without measurable effects on dMSNs properties, M4 receptor affinity, and/or motor behavioral performance^17,32,33^. To overexpress GIRK2 channels in dMSNs, a Cre-dependent virus encoding for GIRK2 and tdTomato fluorophore (AAV.DIO.GIRK2.T2A.tdTomato) was unilaterally injected into the DSt of D1-Cre (*Drd1-Cre^+/-^*) mice at the same time as either saline or the different doses of 6-OHDA were injected into the MFB (Figure 3A-3B). Targeting the center of DSt ensured similar levels of GIRK2 across DMS and DLS as previously shown^17^. After three weeks to allow for protein expression, acute brain slices were prepared and M4 receptor mediated inhibitory postsynaptic currents (M4-IPSC) in tdTomato^+^ GIRK2^+^ dMSNs were measured when ACh release was evoked from ChIs following a single electrical stimulus (25 µA, 0.5 ms) (Figure 3C). Recordings were conducted in the presence of antagonists blocking glutamate, GABA, DA, and nicotinic receptors in order to isolate muscarinic cholinergic transmission.

**Figure 3:**
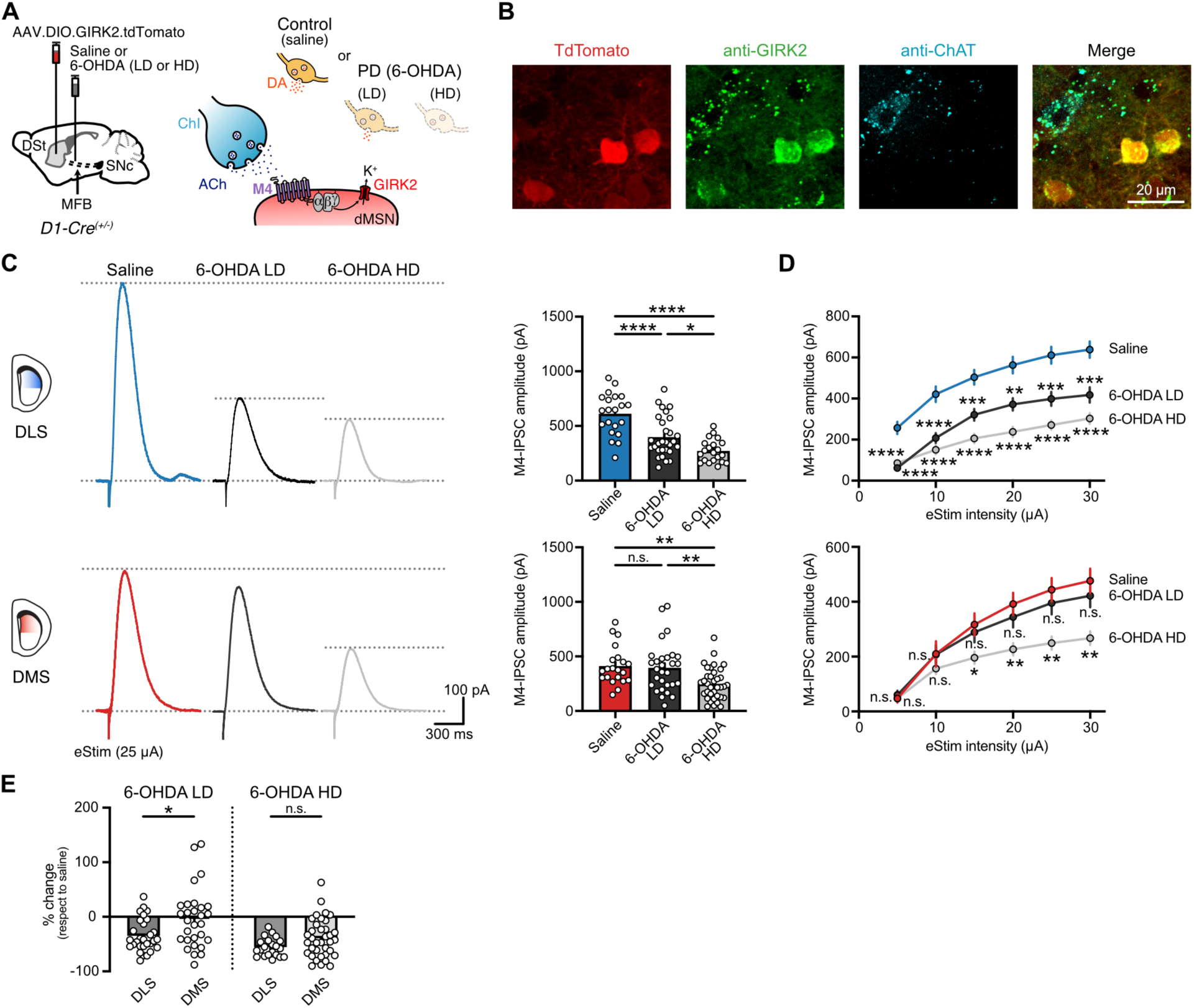
Reductions in M4-signaling are more sensitive to the extent of DA loss in the DLS than in the DMS. **(A)** Schematics of Cre-dependent GIRK2-encoding virus and saline or 6-OHDA (LD or HD) injections into the DSt and MFB from D1-Cre mice respectively (left). Cartoon of ChI-dMSN synapse showing the intracellular coupling between M4 receptor and GIRK2 in dMSNs (right). **(B)** Representative fluorescence image from the DSt showing AAV-induced overexpression of TdTomato and GIRK2 selectively in D1-expressing cells. Note that ChIs (ChAT^+^-cells) express lower levels of endogenous GIRK2. **(C)** Representative traces (left) and quantification (right) of electrically evoked M4-IPSCs (25 µA, 0.5 ms) in DLS (top) and DMS (bottom) for saline, 6-OHDA LD and 6-OHDA HD groups (DLS: Saline n=21, N=6; 6-OHDA LD n=29, N=11; 6-OHDA HD n=21, N=7; One-Way ANOVA p<0.0001; Tukey’s post-hoc test / DMS: Saline n=21, N=11; 6-OHDA LD n=29, N=9; 6-OHDA HD n=36, N=11; Kruskal-Wallis p=0.0007; Dunn’s post-hoc test). **(D)** Plot of M4-IPSCs amplitudes vs electrical stimulation intensity in DLS (top) and DMS (bottom) for saline, 6-OHDA LD and 6-OHDA HD groups (Two-Way RM ANOVA, Treatment: p<0.0001 for DLS and p=0.0067 for DMS, Holm-Šídák’s post-hoc test; only statistical comparisons to saline group are shown). **(E)** Quantification of the percent change in M4-IPSC amplitude following partial or complete DA loss with respect to saline (6-OHDA LD: Mann-Whitney p=0.0181; 6-OHDA HD: unpaired t-test p=0.0533). Summary data is mean ± SEM. Extended statistical data is provided in Supplemental Table S1. n: number of cells, N: number of mice; n.s. p>0.05; *p<0.05; **p<0.01; ***p<0.001; ****p<0.0001.

In the DLS, partial DA loss reduced M4-mediated cholinergic transmission in dMSNs, as could be seen by a decrease (∼35%) in the amplitude of electrically evoked M4-IPSCs from low-dose 6-OHDA-treated mice compared to saline-treated control condition (Figure 3C). This effect was also observed over a range of electrical stimulation intensities (Figure 3D) and depended on the 6-OHDA dose, as the amplitude of M4-IPSCs was further reduced in dMSNs from high-dose 6-OHDA-treated mice (∼55%) (Figure 3C). The decrease in M4-mediated cholinergic transmission in dMSNs also occurred in the DMS (∼40%), but only with a more complete dopaminergic lesion since there was no change in the amplitude of electrically evoked M4-IPSCs in the low-dose 6-OHDA-treated group compared to control condition (Figure 3C). Similar results were observed over a range of electrical stimulation intensities (Figure 3D). Comparing between striatal regions, the partial loss of DA thus selectively impaired M4-transmission in dMSNs in the DLS as it spared the DMS (Figure 3E). Following a nearly complete DA loss, there was a trend towards a more pronounced decrease in the strength of M4-signaling in the DLS (∼55%) than in the DMS (∼40%), but it did not reach significance (Figure 3E). These observations resemble the uneven extent of dopaminergic fiber degeneration across DSt regions, which was observed with the partial lesion but lost when DA depletion was total (Figure 1B, 1D). Thus, the reduction in M4-mediated cholinergic signaling in dMSNs constitutes an adaptation or compensatory change in response to the gradual and progressive dopaminergic degeneration across both striatal regions.

### Reduced M4-signaling is due to a decrease in postsynaptic M4 function across both striatal regions and not due to changes in ACh release or clearance

We previously demonstrated that complete DA loss in the DLS leads to a decrease in postsynaptic M4 receptor function in dMSNs, involving not only downregulation of receptor levels but also impairments in downstream signaling^17^. We next set out to determine whether the same mechanism underlies the decrease in the strength of M4-mediated cholinergic transmission in the DLS following a partial DA loss as well as in the DMS in response to a more complete dopaminergic lesion. To directly examine postsynaptic M4 receptors, we bath applied a saturating concentration of the muscarinic agonist oxotremorine (Oxo-M, 10 µM) and recorded evoked M4-mediated outward currents from dMSNs overexpressing GIRK2 (Figure 4A). Maximum outward currents in the DLS were significantly decreased to a similar extent (∼40%) by both partial and complete dopaminergic lesions (Figure 4B). In contrast, Oxo-M currents in the DMS were only reduced following near total DA depletion and were unaffected when dopaminergic degeneration was moderate (Figure 4B). The decrease was similar for both DSt regions (∼40-45%) (Figure 4C). Together, this indicates that the region-specific reductions in the amplitude of electrically evoked M4-IPSCs driven by DA loss are in all cases due to a comparable decrease in postsynaptic M4 receptor function in dMSNs.

**Figure 4:**
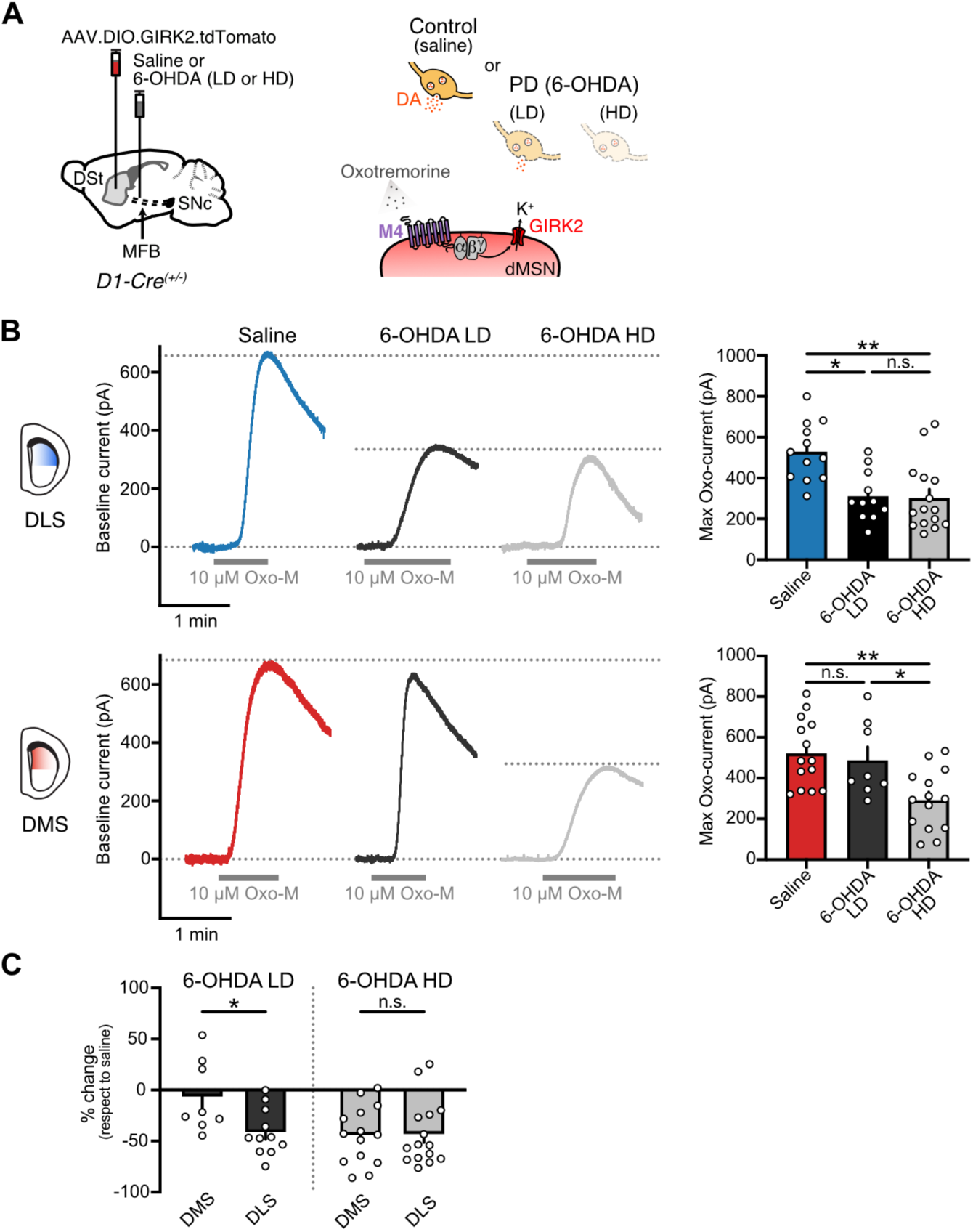
Reduction in M4-transmission due to DA loss is the result of reduced postsynaptic M4 receptor function. **(A)** Schematics of AAV9.hSyn.DIO.tdTomato.T2A.GIRK2 and saline or 6-OHDA (LD or HD) injections into the DSt and MFB respectively in D1-Cre mice (left). Cartoon showing the bath application of muscarinic agonist Oxo-M (right). **(B)** Representative traces of M4 receptor mediated outward currents evoked by bath application of Oxo-M (10 µM) in DLS (top) and DMS (bottom) for saline, 6-OHDA LD and 6-OHDA HD groups (left). Spontaneous M4-IPSCs and electrical artifacts were blanked for clarity. Quantification of maximal Oxo-M currents is shown on the bar graphs (right) (DLS: Saline n=12, N=6; 6-OHDA LD n=11, N=10; 6-OHDA HD n=15, N=7; Kruskal-Wallis p=0.0016; Dunn’s post-hoc test / DMS: Saline n=14, N=10; 6-OHDA LD n=8, N=6; 6-OHDA HD n=14, N=11; One-Way ANOVA p=0.002; Tukey’s post-hoc test). **(C)** Quantification of the percent change in maximal Oxo-M outward currents following partial or complete DA loss with respect to saline (6-OHDA LD: unpaired t-test p=0.0192; 6-OHDA HD: Mann-Whitney p=0.9486). Summary data is mean ± SEM. Extended statistical data is provided in Supplemental Table S1. n: number of cells, N: number of mice; n.s. p>0.05; *p<0.05; **p<0.01; ***p<0.001; ****p<0.0001.

Next, we investigated whether changes in the presynaptic release of ACh and/or its clearance by acetylcholinesterase (AChE) also contribute to the reduction in the strength of M4-mediated cholinergic transmission. Following DA loss, although different alterations in ACh levels have been reported, increases have been canonically ascribed to reduced D2-mediated inhibition of ChI firing^34,35^. As the adaptation in M4-mediated cholinergic transmission in dMSNs was studied while pharmacologically isolating muscarinic receptors, we searched for contributing alterations in ACh release and hydrolysis under these same conditions. Thus, recordings were similarly done in the presence of antagonists for nicotinic, glutamate, GABA and DA receptors to allow us to bypass any additional changes in ChIs that may arise following DA loss. In order to determine the extent of ACh release, we used the genetically encoded optical sensor GRAB_ACh3.0_^36^ and 2-photon (2P) fluorescent imaging (Figure 5A). First, we verified sensor expression levels and examined the dynamic range of the sensor. A virus encoding for GRAB_ACh3.0_ was unilaterally injected into the DSt of WT mice and three weeks following, brain slices were prepared and a saturating concentration of ACh (100 µM) together with the acetylcholinesterase (AChE) inhibitor ambenonium (100 nM) were bath applied (Figure 5A-5B). As there was no difference in ACh-evoked maximal fluorescence between DLS and DMS, similar levels of sensor were present across striatal regions (Figure 5B). Subsequent measurements were always kept below that maximal peak change in fluorescence to ensure that the sensor was not saturated under our experimental conditions.

**Figure 5:**
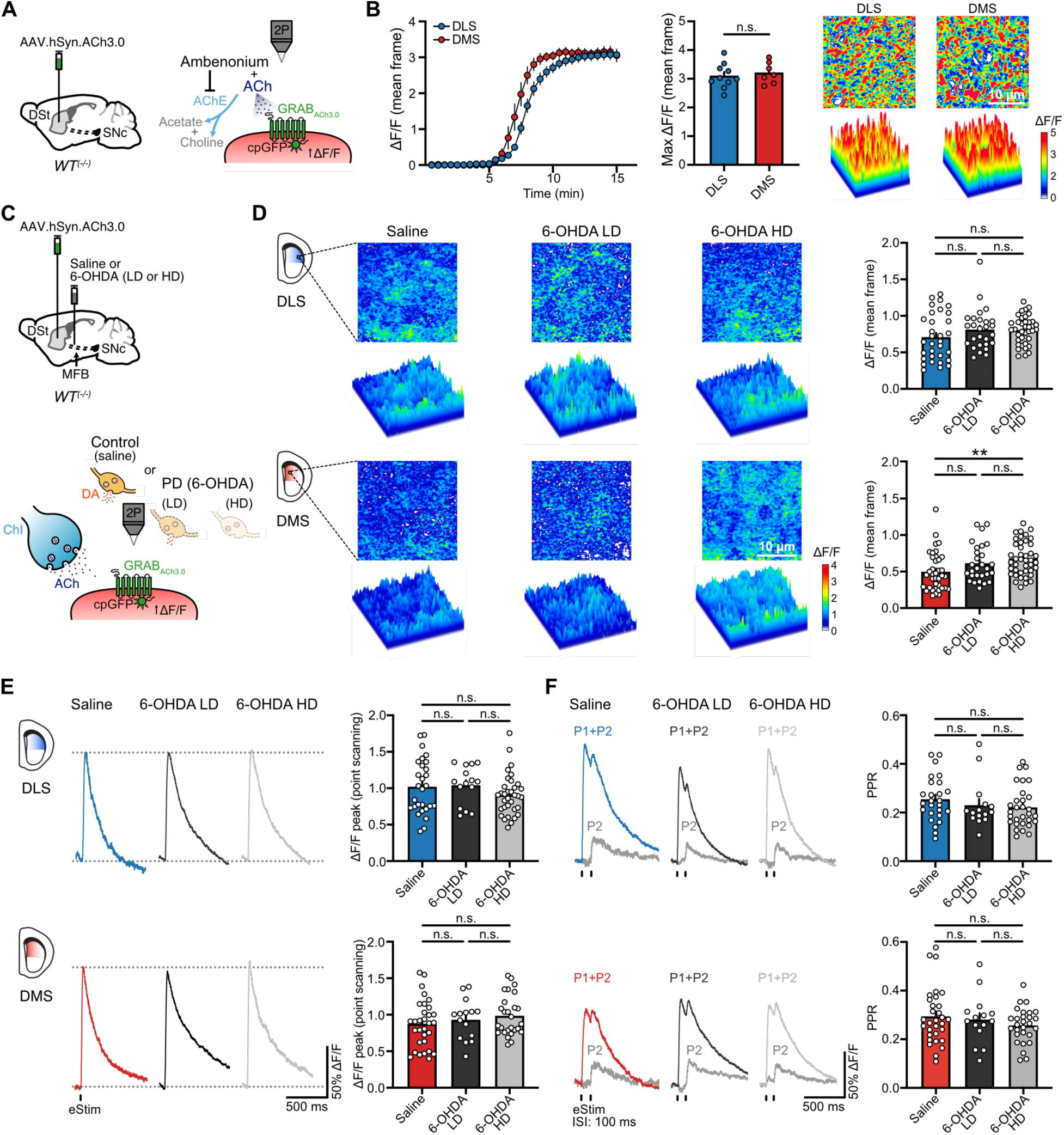
Alterations in presynaptic ACh release in response to DA loss do not contribute to the adaptive reduction in M4-signaling in dMSNs. **(A)** Schematics of AAV9.hSyn.ACh3.0 injection into the DSt of WT mice (left). Cartoon schematic showing the principle of GRAB_ACh3.0_ sensor function and the bath application of ACh with ambenonium (right). **(B)** Time courses of GRAB_ACh3.0_ fluorescence change (mean frame) following bath application of 100 µM ACh and 100 nM ambenonium for DLS and DMS (left). Summary data for maximal ΔF/F (center). Representative images and surface plots showing the maximal changes in GRAB_ACh3.0_ fluorescence across striatal regions (right) (DLS n=10, N=5; DMS n=7, N=4; unpaired t-test p=0.5967). **(C)** Schematics of AAV9.hSyn.ACh3.0 and saline or 6-OHDA (LD or HD) injections into the DSt and MFB respectively in WT mice (top). Cartoon depicting how ACh released from ChIs is detected by GRAB_ACh3.0_ expressed in striatal cells (bottom). **(D)** Representative images and surface plots of GRAB_ACh3.0_ fluorescence changes (mean frame) in DLS (top) and DMS (bottom) following application of a single electrical stimulus (25 µA, 0.5 ms) in the center of the striatal region of interest (left). Quantification of ΔF/F is shown on the bar charts (right) (DLS: Saline n=32, N=9; 6-OHDA LD n=27, N=6; 6-OHDA HD n=38, N=9; Kruskal-Wallis p=0.2419 / DMS: Saline n=38, N=9; 6-OHDA LD n=28, N=6; 6-OHDA HD n=42, N=9; Kruskal-Wallis p=0.0019; Dunn’s post-hoc test). **(E)** Representative 2P-point scanning traces and quantification of fluorescence changes of GRAB_ACh3.0_ (DLS: Saline n=28, N=6; 6-OHDA LD n=15, N=4; 6-OHDA HD n=34, N=5; Kruskal-Wallis p=0.1903 / DMS: Saline n=31, N=7; 6-OHDA LD n=15, N=4; 6-OHDA HD n=27, N=5; Kruskal-Wallis p=0.467). **(F)** Representative 2P-point scanning traces and summary data of GRAB_ACh3.0_ fluorescence changes following paired-pulse stimulation (P1+P2), with the digitally subtracted P2 component shown on gray (DLS: Saline n=26, N=6; 6-OHDA LD n=15, N=4; 6-OHDA HD n=30, N=5; Kruskal-Wallis p=0.2159 / DMS: Saline n=31, N=7; 6-OHDA LD n=15, N=4; 6-OHDA HD n=28, N=5; Kruskal-Wallis p=0.5913). Summary data is mean ± SEM. Extended statistical data is provided in Supplemental Table S1. n: number of cells, N: number of mice; n.s. p>0.05; *p<0.05; **p<0.01; ***p<0.001; ****p<0.0001.

To examine changes in parkinsonian mice, the virus encoding for GRAB_ACh3.0_ was then unilaterally injected into the DSt of WT mice at the same time as either of the different doses of 6-OHDA or saline were injected into the MFB (Figure 5C). Three weeks following, brain slices were prepared and a single electrical stimulation (25 µA, 0.5 ms) was used to evoke the synaptic release of ACh as before, while 2P imaging was performed to measure changes in GRAB_ACh3.0_ fluorescence in frames from square regions of interest (25µm x 25µm) (Figure 5D). Neither a partial nor total DA depletion had an effect on ACh release in the DLS (Figure 5D). In the DMS, while ACh release was unaffected when the moderate lesion was induced by the low-dose of 6-OHDA, it was slightly increased following the near complete loss of DA produced by the high-dose of 6-OHDA (Figure 5D). As frame-scans of regions of interest have relatively slow temporal resolution, we also performed high speed 2P point scanning to better resolve peaks in ACh release at selected “hotspots” (Figure 5E). The maximum peak in GRAB_ACh3.0_ fluorescence using this approach was similar between control and 6-OHDA-treated animals not only in the DLS, but also in the DMS (Figure 5E). We also found that the paired-pulse ratio (PPR) following two stimulations (inter stimulus interval (ISI): 100ms) was unaffected by either low- or high-dose of 6-OHDA treatments in both striatal regions, indicating that the probability of release was also not altered under these conditions (Figure 5F). Altogether, these results suggest that, under the present conditions, changes in the presynaptic release of ACh do not contribute to the decrease in the strength of M4-mediated cholinergic transmission in dMSNs.

As ACh levels are regulated by AChE-mediated hydrolysis, we next tested if enzymatic clearance was affected following partial or complete DA lesions, thus contributing to reduced M4-signaling. To examine this, we bath applied the AChE inhibitor ambenonium (10 nM) while again recording electrically-evoked M4-IPSCs in GIRK2-expressing dMSNs (Figure 6A). As expected, the net charge transfer of evoked synaptic currents was significantly increased in all cases following AChE inhibition (Figure 6B). However, there was no difference in the effect of ambenonium across DMS or DLS after treatment with either the low- or the high-dose of 6-OHDA (Figure 6B). Together, these results suggest that changes in evoked ACh release from ChIs and/or ACh clearance by AChE do not directly contribute to the reduction in M4-mediated cholinergic transmission in dMSNs, further supporting the decrease in postsynaptic M4 receptor function as the sole mechanism underlying this adaptation in the direct pathway in response to dopaminergic lesions.

**Figure 6:**
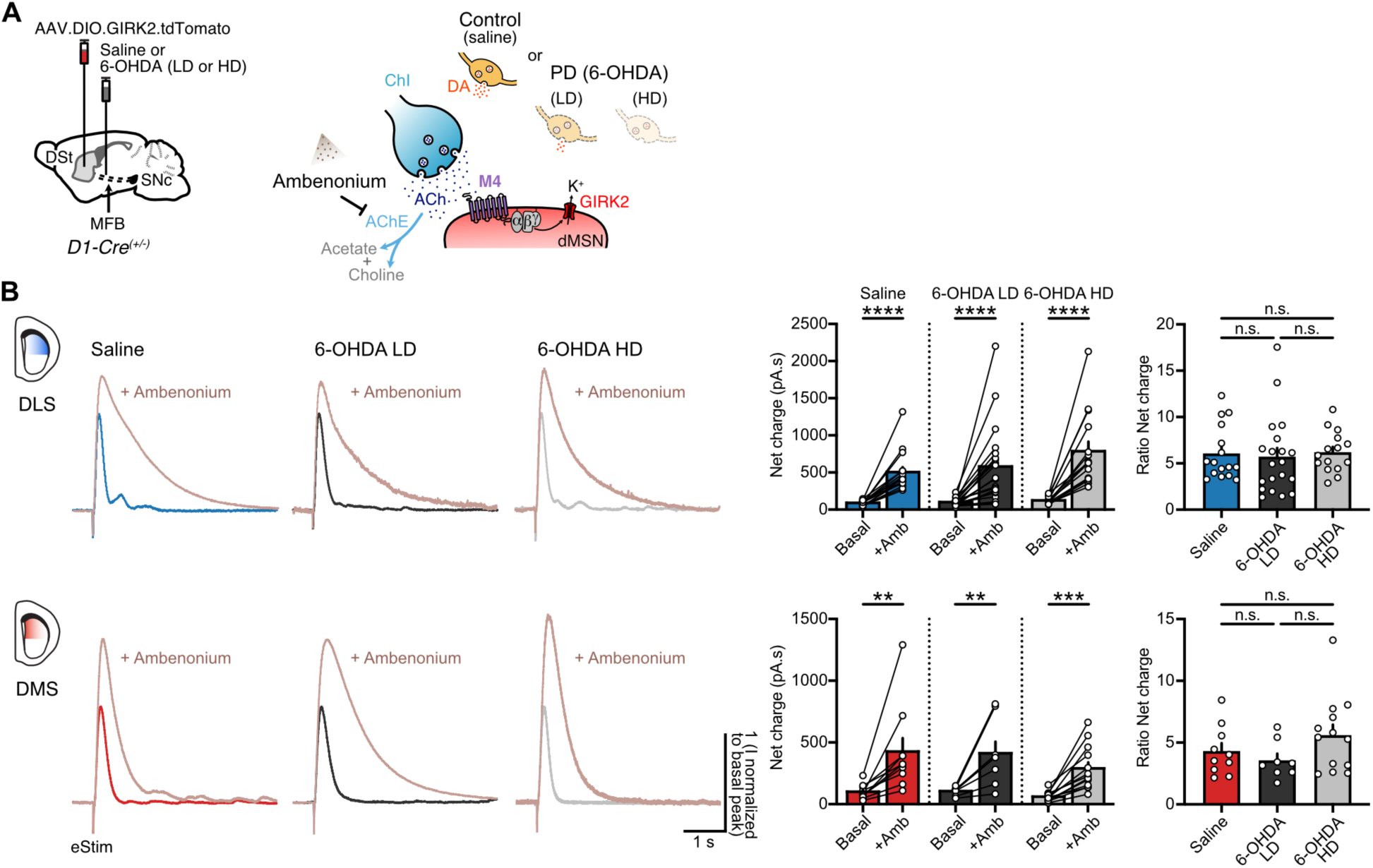
ACh clearance by AChE across striatal regions remains unaffected following either partial or complete DA loss. **(A)** Schematics of Cre-dependent GIRK2-encoding virus and saline or 6-OHDA (LD or HD) injections into the DSt and MFB from D1-Cre mice respectively (left). Cartoon of ChI-dMSN synapse depicting the bath application of ambenonium to block ACh hydrolysis by AChE (right). **(B)** Representative traces of electrically evoked M4-IPSCs (left) before (Basal) and after (+Amb) bath application of ambenonium (10 nM) in DLS (top) and DMS (bottom). Quantification of net charge as absolute values (center) and normalized to control (right) (DLS: Saline n=16, N=4; 6-OHDA LD n=20, N=8; 6-OHDA HD n=15, N=5; Wilcoxon tests p<0.0001 for absolute values and Kruskal-Wallis p=0.4736 for ratio / DMS: Saline n=10, N=6; 6-OHDA LD n=8, N=6; 6-OHDA HD n=13, N=8; Wilcoxon test p=0.002 for saline, paired t-test p=0.0082 for 6-OHDA LD and paired t-test p=0.0003 for 6-OHDA HD in absolute values and One-Way ANOVA p=0.1687 for ratio). Summary data is mean ± SEM. Extended statistical data is provided in Supplemental Table S1. n: number of cells, N: number of mice; n.s. p>0.05; *p<0.05; **p<0.01; ***p<0.001; ****p<0.0001.

### Reduced M4-mediated cholinergic signaling cannot be reversed by L-DOPA administration even when DA is partially lost

Prolonged replacement therapy with the DA precursor L-DOPA is the most efficacious treatment for PD especially at early stages, but can lead to levodopa-induced dyskinesia (LID) as the disease progresses^3^. We previously showed that chronic L-DOPA treatment that led to LID in parkinsonian mice completely depleted of DA by a high-dose of 6-OHDA could not revert the impairment in M4-signaling in dMSNs in the DLS^17^. We next wanted to determine whether L-DOPA treatment was instead able to reverse the reduction in M4 receptor function following partial DA lesions, mimicking earlier stages of degeneration when LID has not yet developed. To test this, low-dose 6-OHDA-lesioned mice overexpressing GIRK2 in dMSNs were treated for 7-8 days with daily i.p. injections of L-DOPA (2 mg/kg) plus benserazide (12 mg/kg) (Figure 7A). Acute brain slices were prepared 30 min – 1 h after the last L-DOPA administration and M4-mediated cholinergic transmission in GIRK2^+^ dMSNs was examined (Figure 7A). The amplitude of electrically evoked M4-IPSCs in the DLS from partially lesioned mice treated with L-DOPA was still significantly reduced compared to saline-treated animals, while there were no changes when compared to the low-dose 6-OHDA-treated mice (Figure 7B). In the DMS, and as expected since there were no changes in the strength of M4-signaling following a moderate loss of DA, the amplitude of the M4-IPSCs was also unaffected (Figure 7B).

**Figure 7:**
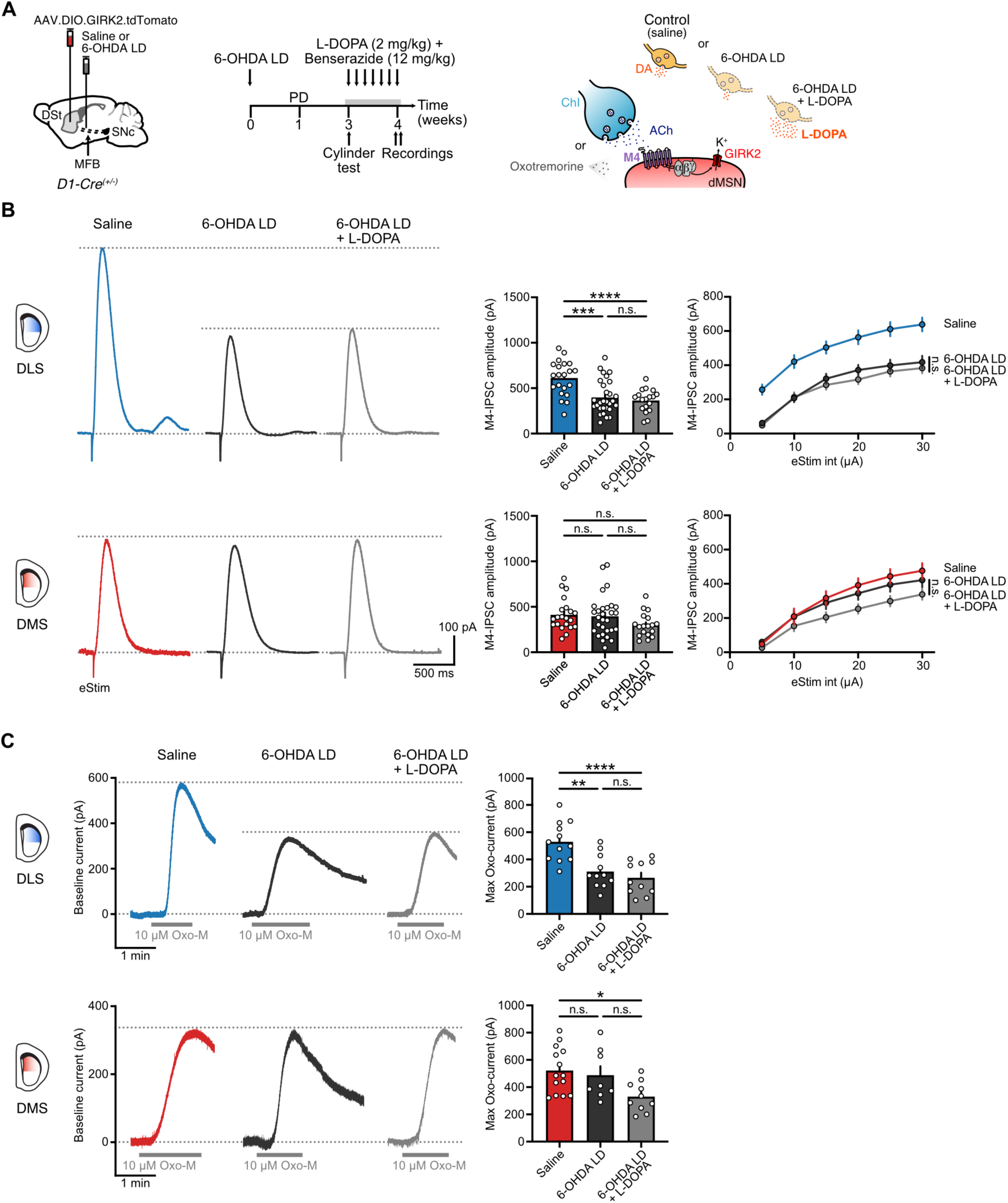
L-DOPA does not revert the impairment in M4-signaling following partial DA loss. **(A)** Schematics of Cre-dependent GIRK2-encoding virus and saline or 6-OHDA LD injections into the DSt and MFB from D1-Cre mice respectively (left). Timeline for partial DA depletion, prolonged L-DOPA treatment and experiments (center). Cartoon of ChI-dMSN synapse depicting the experimental conditions (right). **(B)** Representative traces (left) and quantification (right) of electrically evoked M4-IPSCs (25 µA, 0.5 ms) in DLS (top) and DMS (bottom) for saline, 6-OHDA LD and 6-OHDA LD + L-DOPA groups. Data from saline and 6-OHDA LD conditions is taken from Figure 3C (DLS: Saline n=21, N=6; 6-OHDA LD n=29, N=11; 6-OHDA LD + L-DOPA n=18, N=4; One-Way ANOVA p<0.0001; Tukey’s post-hoc test / DMS: Saline n=21, N=11; 6-OHDA LD n=29, N=9; 6-OHDA LD + L-DOPA n=18, N=4; Kruskal-Wallis p=0.1197). **(C)** Plot of M4-IPSCs amplitudes vs electrical stimulation intensity in DLS (top) and DMS (bottom) for saline, 6-OHDA LD and 6-OHDA LD + L-DOPA groups. Data from saline and 6-OHDA LD conditions is taken from Figure 3D (Two-Way RM ANOVA Treatment: p<0.0001 for DLS and p=0.1261 for DMS; Holm-Šídák’s post-hoc test, only statistical comparisons between 6-OHDA LD and 6-OHDA LD + L-DOPA groups are shown). **(D)** Representative traces of M4 receptor mediated outward currents evoked by bath application of Oxo-M (10 µM) in DLS (top) and DMS (bottom) for saline, 6-OHDA LD and 6-OHDA LD + L-DOPA groups (left). Spontaneous M4-IPSCs and electrical artifacts were blanked for clarity. Quantification of maximal Oxo-M currents is shown on the bar graphs (right). Data from saline and 6-OHDA LD conditions is taken from Figure 4B (DLS: Saline n=12, N=6; 6-OHDA LD n=11, N=10; 6-OHDA LD + L-DOPA n=11, N=4; One-Way ANOVA p<0.0001; Tukey’s post-hoc test / DMS: Saline n=14, N=10; 6-OHDA LD n=8, N=6; 6-OHDA LD + L-DOPA n=10, N=4; One-Way ANOVA p=0.0191; Tukey’s post-hoc test). Summary data is mean ± SEM. Extended statistical data is provided in Supplemental Table S1. n: number of cells, N: number of mice; n.s. p>0.05; *p<0.05; **p<0.01; ***p<0.001; ****p<0.0001.

Next, we directly examined postsynaptic M4 receptor function by bath applying a saturating concentration of Oxo-M as previously described. Similar to our observation of M4-IPSCs, maximum outward currents in the DLS were still significantly reduced following a partial depletion of DA despite L-DOPA treatment, being no different from the low-dose 6-OHDA-treated condition (Figure 7C). In the DMS, maximal M4-mediated outward currents also remained unchanged (Figure 7C). Together, these results suggest that the adaptive or compensatory reduction in postsynaptic M4 receptor function in dMSNs driven by the loss of endogenous DA cannot be rescued by replacement with exogenous L-DOPA despite the presence of surviving dopaminergic fibers in partially lesioned animals.

## DISCUSSION

Similar to the many adaptive or compensatory changes in striatal circuits that arise in response to DA loss, the reduction in M4-mediated cholinergic signaling was originally identified in DLS dMSNs of mice completely lacking dopaminergic fibers mimicking advanced stages of PD^17^. However, whether this alteration initiates at earlier stages in the DLS and/or also occurs at the DMS has remained unexplored. In the present work, we found that the decrease in direct pathway cholinergic modulation through M4 receptors was driven by DA loss, since it closely followed the temporal and spatial pattern of dopaminergic degeneration. This adaptive change resulted from reduced postsynaptic M4 receptor function, starting in the DLS following a partial decline in striatal DA and then progressing to the DMS with more complete lesions. Even at early stages of degeneration, replacing DA by administering its precursor L-DOPA failed to rescue the impairment in M4-signaling. Due to the therapeutic value of targeting M4 receptors^37,38^ understanding how direct pathway cholinergic modulation is progressively altered as DA declines over time and across striatal regions provides key information to comprehend its contribution to motor and cognitive symptoms of PD.

PD is characterized by the progressive loss of DA in time and space as degeneration initiates in the putamen or DLS at earlier stages^19,20,39,40^, leading to motor dysfunction, and then proceeds to the caudate or DMS at more advances stages, where it contributes to neuropsychiatric manifestations such as cognitive decline and depression^41,42^. As motor symptoms do not appear until striatal DA innervation has decreased by 50-70%, compensatory mechanisms are thought to contribute to the maintenance of motor performance despite strong DA loss^19,20,22,43^. Most compensations rely on changes in dopaminergic neurons to elevate extracellular DA levels, but additional adaptations in striatal and extrastriatal non-dopaminergic systems aiming to reduce the imbalance between striatal output pathways have also been reported^44–46^. Exploring homeostatic mechanisms in PD patients at early stages, however, has been hindered by the difficulty of conducting longitudinal studies and the late time point of diagnosis, thus requiring the use of parkinsonian animal models to study progressive aspects of DA decline. We found that when inducing a partial dopaminergic lesion with a low-dose of 6-OHDA, M4-signaling in dMSNs was reduced only in the DLS. This was in contrast to the DMS where M4-signaling impairments occurred only following the total depletion of DA with a high-dose of 6-OHDA. Thus, the adaptive decrease in M4-mediated cholinergic modulation of dMSNs closely parallels or resembles the uneven pattern of progressive dopaminergic degeneration over time and across striatal regions.

For both partial and complete DA depletions, we found that the reduced strength of M4-transmission in dMSNs was due to decreases in M4 receptor function, since ACh release and clearance were largely unaffected. Although we did observe a slight increase in ACh release in the DMS following complete dopaminergic lesion, this increase cannot explain a reduction in M4-signaling. Instead, it might explain the trend towards a less pronounced decrease in the DMS compared to the DLS despite the similar reduction in postsynaptic M4 receptor function. In the DMS, the primary mechanism of dopaminergic modulation of ChI firing is thought to be D2-mediated inhibition^32,47–49^, which remains relatively conserved when DA is partially depleted due to dopaminergic homeostatic mechanisms^29^. Although these compensations are likely overwhelmed at advanced stages of degeneration, potentially resulting in higher ACh release when ChI pauses in firing are reduced or lost^10^, all our experiments were conducted in the presence of a D2 receptor antagonist. Interestingly, the increase in release was detected only when measuring the change in sensor fluorescence as the average of a frame, but not when the peak was determined at higher temporal and spatial resolution by 2P point scanning. As PPR was also unaffected, this might suggest an increase in the area where ACh was released instead of an increase at individual release sites. While an augmented availability of ACh due to ChI axonal overgrowth in response to DA loss has been proposed^6^ and a recent work has reported an increase in the binding of vesicular ACh transporter (VAChT) in PD patients^16^, most morphological studies in the 6-OHDA model have focused on alterations in the ChI dendritic tree^15,50^. Further work is thus needed to compare potential differences in ChI axonal arborizations across DSt regions following DA depletion.

In our previous work we showed that reduced postsynaptic M4 function in DLS dMSNs is due to a downregulation in M4 receptor levels and downstream signaling, with regulator of G-protein signaling 4 (RGS4) playing a key role^17^. DA loss could drive this cholinergic adaptation directly in the dMSNs and/or indirectly, through changes in excitability of other striatal cells and transmitter release. In the first case, the reduced levels of cAMP following the diminished Gα_olf/s_-coupled D1 receptor signaling could negatively regulate expression and activity of Gα_i/o_-coupled receptors such as M4 receptor^51^ and even increase the transcription of regulatory proteins like RGS4^11,52–54^. Moreover, DA depletion has been shown to directly increase dMSN excitability as a homeostatic response^13,55,56^ and reducing inhibitory M4-signaling might add to this mechanism. In the second scenario, indirect effects might arise from DA-depletion-induced changes in ChI excitability and ACh tonic and/or phasic release. Although conflicting results have been reported^6,7,9–15^, the prevailing view postulates that PD is characterized by hyperexcitable ChIs^12^ and increased ACh levels^9,57^. This might lead to a decrease in M4 receptor levels by agonist-induced internalization, subcellular redistribution and degradation^58,59^. Future studies are therefore needed to examine whether the *in vitro* decreases in dMSN postsynaptic M4 receptor function translate into reduced M4-mediated cholinergic modulation of the direct pathway *in vivo*, where other compensatory and pathological alterations driven by DA loss may act in concert to alter striatal output and behavior.

Understanding the temporal progression of non-dopaminergic alterations in PD is key for developing alternative and dynamic therapeutics approaches that reduce reliance on L-DOPA, decrease the incidence of LID, and alleviate DA unresponsive symptoms in the long-term. Here we showed that L-DOPA administration was unable to revert the adaptive reduction in dMSNs M4-signaling when mimicking early stages of dopaminergic degeneration. This suggests that it is unlikely that non-dopaminergic alterations in response to DA loss can be rescued at the time motor symptoms arise and the disease is diagnosed^22^, and even less likely when L-DOPA treatment is initiated, which is usually further delayed from diagnosis^60,61^. However, it has been shown that L-DOPA administration at early stages of degeneration can normalize vesicular levels of DA in surviving dopaminergic fibers and partially restore physiological patterns of signaling^62,63^, which might help to prevent, delay and/or counteract the changes in non-dopaminergic systems that develop in response to the loss of endogenous DA. Future studies in mouse models with even more moderate DA lesions will be required to precisely determine when cholinergic adaptations start as well as the potential of L-DOPA to revert them at the different disease stages.

While the present work focused primarily on the alterations resulting from DA loss, we also noted regional differences in M4 receptor signaling that exist in the intact or non-lesioned DSt. We found that the strength of transmission was slightly greater in the DLS compared to the DMS (Figure 3), which was likely due to differences in the extent of ACh released across regions (Figure 5) since postsynaptic M4 receptor function was similar in the DLS and DMS (Figure 4). The greater density of ChAT^+^ neuropil in the DLS^64,65^, a greater number of ACh release sites, or differences in modulatory drive onto ChIs by metabotropic receptors (mGluRs)^32,47,48^ may potentially account for the differential ACh release across regions. Future work exploring regional differences in non-dopaminergic systems that also modulate striatal output, such as the ones uncovered here for the cholinergic system, will be critical for providing insight into how the distinctive behaviors associated to each striatal subdivision are shaped.

Together, our results demonstrate that a reduction in M4-mediated cholinergic modulation of the striatal direct pathway develops in response to DA loss parallelling the temporal and regional pattern of degeneration in a mouse model mimicking early and advanced stages of PD. Characterizing the progression of non-dopaminergic adaptive changes over time and across regions, especially for ACh which closely interacts with DA, is paramount for understanding PD and dynamically targeting its symptoms according to the disease stage.

## Data and Materials Availability

All generated data is available in the main text, Supplemental Table S1 or at Zenodo repository (https://doi.org/10.5281/zenodo.14956884)

The data, code, protocols, and key lab materials used in this study are listed in a Key Resource Table alongside their persistent identifiers at Zenodo repository (https://doi.org/10.5281/zenodo.14956884)

No code was generated for this study; all data cleaning, preprocessing, analysis, and visualization was performed using commercial software stated in Materials and Methods.

## Author contributions

BEN and CPF designed experiments. BEN performed research experiments and analyzed the data. BEN and CPF wrote the manuscript.

## Acknowledgments

This work was funded by NIH grants R01-NS95809 (CPF), R01-DA35821 (CPF), Parkinson’s Foundation Postdoctoral Fellowship for Basic Scientists PF-PRF-839073 (BEN), as well as funded in part by Aligning Science Across Parkinson’s (ASAP-020529) (CPF) through the Michael J. Fox Foundation for Parkinson’s Research (MJFF). For the purpose of open access, the author has applied a CC BY public copyright license to all Author Accepted Manuscripts arising from this submission.

---

**Supplemental Table S1.**
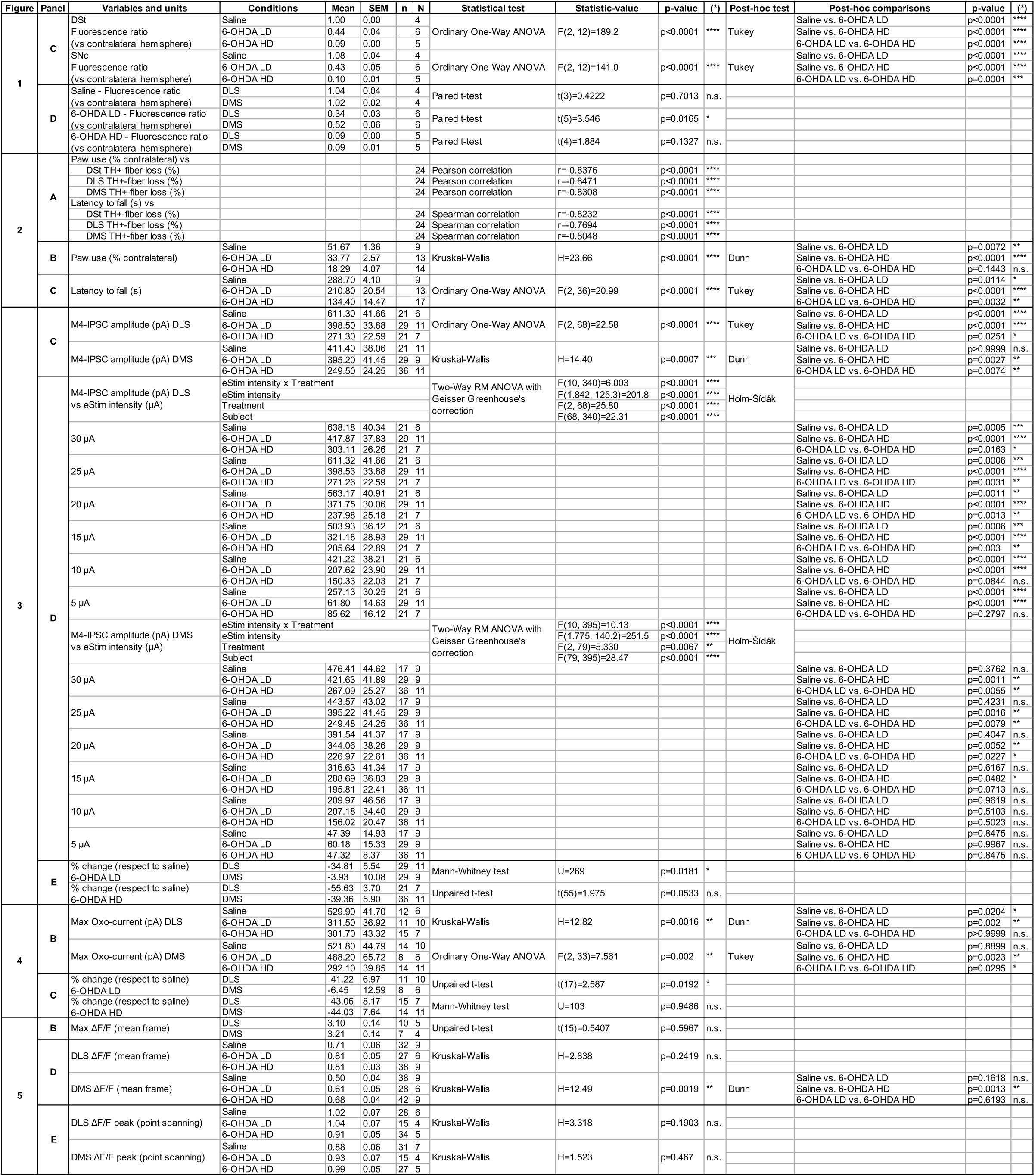
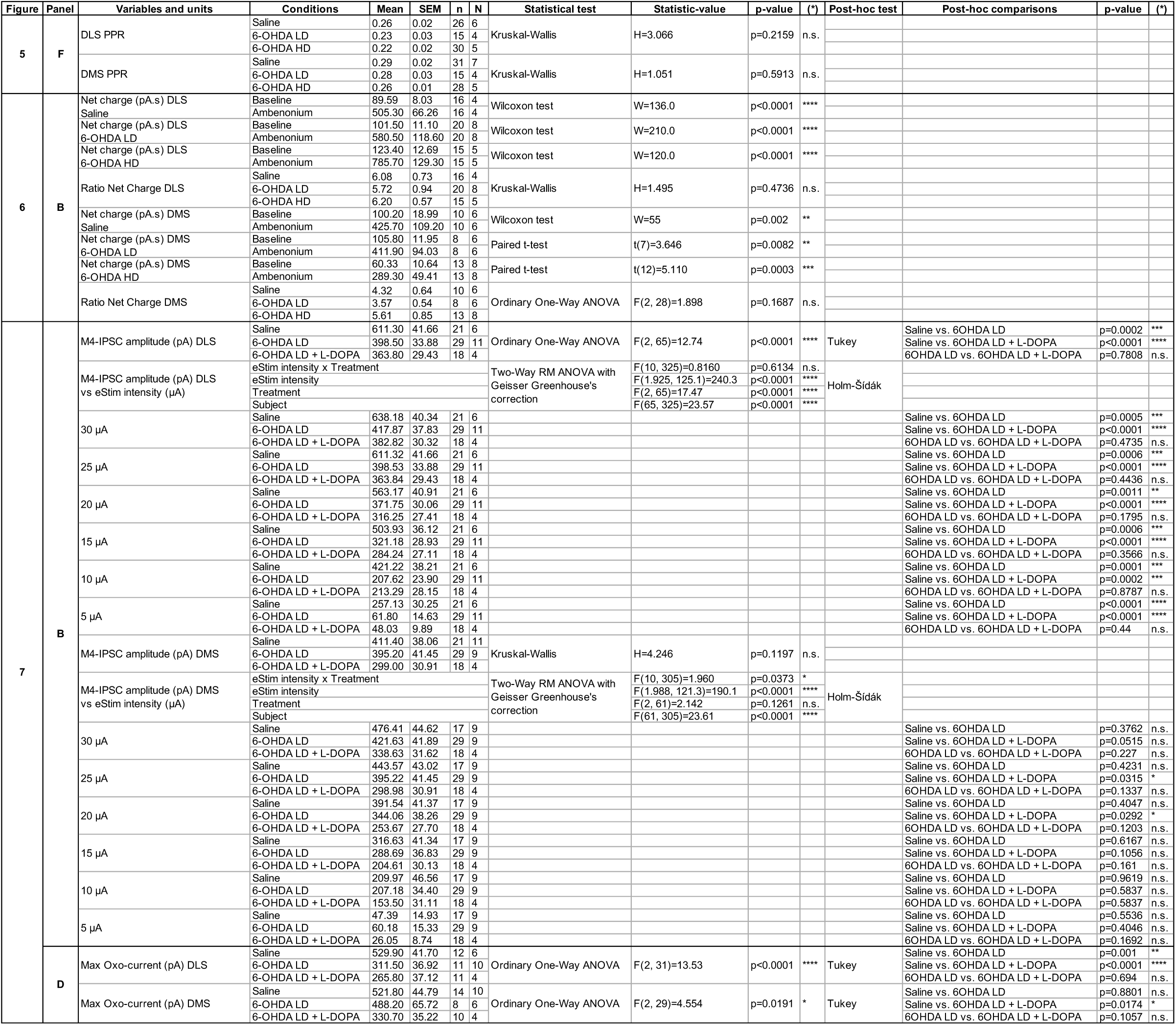
Extended statistical data.

